# Accelerated pseudogenization in ancient endosymbionts of giant scale insects

**DOI:** 10.1101/2024.10.23.619753

**Authors:** Jinyeong Choi, Pradeep Palanichamy, Hirotaka Tanaka, Takumasa Kondo, Matthew E. Gruwell, Filip Husnik

## Abstract

Symbiotic microorganisms are subject to a complex interplay of environmental and population-genetic pressures that drive their gene loss. Despite the widely held perception that ancient symbionts have stable genomes, even tiny genomes experience ongoing pseudogenization. Whether these tiny genomes also experience bursts of rapid gene loss is, however, less understood. Giant scale insects (Monophlebidae) feed on plant sap and rely on the symbiotic bacterium *Walczuchella* which provides them with essential nutrients. When compared to other ancient symbionts with similar genome sizes such as *Karelsulcia, Walczuchella*’s genome was previously reported as unusually pseudogene-rich (10 % of coding sequences). However, this result was based on only one genome assembly raising questions about the assembly quality or a recent ecological shift such as co-symbiont acquisition driving the gene loss. Here, we generated six complete genomes of *Walczuchella* from three genera of giant scales, each with distinct co-symbiotic partners. We show that all the genomes are highly degraded and particularly genes related to the cellular envelope and energy metabolism seem to be ongoing pseudogenization. Apart from general mechanisms driving genome reduction such as the long-term intracellular lifestyle with transmission bottlenecks, we hypothesize that a more profound loss of DNA replication and repair genes together with recent co-obligate symbiont acquisitions likely contribute to the accelerated degradation of *Walczuchella* genomes. Our results highlight that even ancient symbionts with small genomes can experience significant bursts of gene loss when stochastic processes erase a gene that accelerates gene loss or when the selection pressure changes such as after cosymbiont acquisition.

## Introduction

Genome erosion is a hallmark of symbiotic microorganisms [1]. Its outcome depends on an interplay of multiple evolutionary forces affecting host-restricted populations that are bottlenecked every generation [2]. Under relaxed purifying selection, deleterious mutations are more likely to occur and can become easily fixed in populations with small effective sizes and accumulate through Muller’s ratchet effect on asexual populations [2, 3]. The enhanced mutation rate is highly associated with prokaryote genome reduction [4] because the persistent accumulation of nucleotide substitutions disrupts the open reading frames of genes under relaxed selective constraints and eventually leads to the loss of their function (pseudogenization) [5, 6]. For instance, the genomes of recently evolved symbionts can contain over 50% of pseudogenes [7]. Moreover, the reduction of DNA replication and repair genes in symbionts can further accelerate their pseudogenization and overall genome degradation [7].

Intracellular symbionts that are maternally transmitted often contain extremely small genomes [1]. These ancient symbionts have been often maintained in their hosts for hundreds of millions of years due to their crucial role in synthesizing essential nutrients such as amino acids and B vitamins necessary for host development and reproduction [1]. Despite their minimal gene sets, some ancient nutritional symbionts have been reported to experience an ongoing loss of essential genes [1]. In hosts with degraded ancient symbionts, co-occurring microbial partners often complement important metabolic pathways of the original symbionts. Such multipartite symbiotic consortia have been observed in various plant sap-feeding insects, for example, adelgids [8], aphids [9], mealybugs [10, 11], and Auchenorrhyncha [12, 13]. However, the establishment of these new partnerships may induce additional degradation of the ancient symbiont due to the availability of metabolites (or even proteins) provided by the new partner [9].

The Flavobacteriales are a major group of insect symbionts, some of which have evolved long-term relationships with their hosts. Examples of these ancient symbionts include *Candidatus* Karelsulcia in Auchenorrhynchan insects and *Blattabacterium* in cockroaches and *Mastotermes* termites, which play important roles in nutrient provisioning and nitrogen recycling [14, 15]. Other Flavobacteriales, such as *Candidatus* Skilesia from *Geopemphigus* aphids, and *Candidatus* Shikimatogenerans and *Candidatus* Bostrichicola from bostrichid beetles, are associated with nutritional provisioning or cuticle hardness [16, 17]. Many species of scale insects also house symbiotic Flavobacteriales [18–23]. However, the flavobacterial symbionts from different scale insect families form distinct clades that are incongruent with the host phylogeny, suggesting multiple acquisitions and replacements of different symbionts during scale insect evolution [20, 23–25]. However, only two genomes of *Candidatus* Walczuchella and *Candidatus* Uzinuria have been analyzed so far from these diverse flavobacterial symbionts of scale insects [26, 27].

Monophlebidae, also known as giant scales, is a group of scale insects comprised of 267 species in 48 genera [28]. The family, for example, includes the enigmatic cottony cushion scale, *Icerya purchasi* Maskell, which is one of a few androdioecious insects and can cause significant damage to a variety of economically important plants, particularly citrus trees [29] [30]. Giant scales have nutritional symbionts, including *Walczuchella* and multiple co-symbionts related to *Cedecea, Sodalis*, and *Wolbachia* [20, 31, 32]. These obligate symbionts are found in the bacteriomes of the host insects and are vertically transmitted to their offspring [27, 31, 32]. The genome of *Walczuchella* is 309 kbp in size and contains 271 protein-coding sequences [27], but it has a higher number of pseudogenes when compared to other anciently established symbionts, which typically have small genomes with relatively few pseudogenes. Interestingly, the reason for the high level of pseudogenization in *Walczuchella* remains unclear. Here, we addressed this question via metagenome sequencing of six species of giant scales and also determined the symbiont composition of these insects.

## Materials & methods

### DNA extraction and genome sequencing

Six species from three genera of Monophlebidae were used for metagenome sequencing (Table S1). All the samples were preserved in 99% of ethanol at –20°C. For DNA extraction, 1–8 individuals of each species were selected under a dissecting microscope. To eliminate any potential surface contaminants, wax secretions were removed, and the specimens were washed with 99% ethanol. The insects were then ground in liquid nitrogen with a mortar and pestle. DNA was extracted using DNeasy Blood and Tissue Kit (Qiagen) and the MasterPure Complete DNA Purification Kit (Epicenter), following the manufacturers’ protocols. DNA quantity and quality were checked with NanoDrop (Thermo Fisher Scientific) and Qubit 4 fluorometer (Invitrogen). PCR-free DNA libraries were prepared with the NEBNext Ultra II kit (NEB). The libraries were multiplexed and sequenced on the Illumina NovaSeq 6000 and MiSeq sequencers at Okinawa Institute of Science and Technology Graduate University and Pennsylvania State University (Table S1). The quality of raw Illumina reads was assessed using FastQC v0.11.7. [33], and low-quality reads and adapters were removed using fastp v.0.20.0. [34].

### Genome assembly and annotation

Raw Illumina reads were assembled using SPAdes v3.15 [35] using multiple k-mers (Table S1). The taxonomic assignment of assembled scaffolds was conducted using megablast search (NCBI-BLAST v.2.11.0) against the NCBI nucleotide database. *Walczuchella* genome scaffolds were extracted based on the taxon-annotated-GC-Coverage plots generated by Blobtools v1.1.1 [36]. The circular genomes of *Walczuchella* were visually assessed with Bandage v.0.9 [37]. The draft genome assemblies were then polished using Pilon v.1.24 [38] and annotated with Prokka v.1.14.6 [39]. Some hypothetical or unannotated proteins identified by Prokka were manually re-annotated through BLASTp searches against the NCBI Ref-Seq database [40]. Coding sequences (CDSs) were sorted into clusters of orthologous groups (COGs) using eggNOG-mapper v2 [41] with default parameters. The transfer RNAs predictions were confirmed using tRNAs-can-SE v2.0.11 [42]. Genome maps were created using DNAPlotter [43]. A synteny plot of *Walczuchella* genomes was produced using Processing3 (https://processing.org/) based on Blast genome alignments and Prokka annotations. The comparative analyses were performed using genome assemblies of other ancient symbionts obtained from NCBI.

### Pseudogene identification and substitution rate estimation

The putative pseudogenes in the seven complete and circularized *Walczuchella* genomes were predicted using PseudoFinder v1.1.0 [44] using DIAMOND BlastP/BlastX searches v.2.0.4.142 [45] against the non-redundant protein (NR) database. As a reference, pseudogenes were also predicted using the same settings in the published genome assemblies of other ancient symbionts. Using the CDSs of the seven genomes of *Walczuchella*, we identified 190 single-copy orthologous genes with OrthoFinder v.2.5.4 [46]. Of these, 181 single-copy orthologous genes shared by all seven genomes were selected for substitution rate and diversifying selection analysis (after excluding pseudogenes). Two different algorithms, CODEML implemented in PAML v.4.9 [47] and BUSTED in HYPHY v.2.5 [48, 49] were used and the results compared. The unrooted tree of the seven *Walczuchella* genomes was generated using RAxML v.8.2.12 [50] with 1,000 bootstrap replicates and the PROTGAMMAGTR model. The orthologous gene sequences were aligned by MAFFT v.7 [51]. For the analysis with CODEML, nucleotide sequences were translated into amino acid sequences using Geneious Prime v.2023.0.4 [52], and then aligned with MUSCLE v.5.1 [53]. The codon-based DNA alignment was generated using PAL2NAL v.14 [54]. The software packages CODEML (M0 model) and BUSTED were used to calculate the average ratio of non-synonymous to synonymous substitutions (ω = *dN*/*dS*) across the whole gene. The BUSTED analysis produced two values of *dN*/*dS*, one based on relative GTR branch lengths and nucleotide substitution biases, and the other based on improved branch lengths, nucleotide substitution biases and a full codon model. Genes with a *dN*/*dS* ratio between 0.95 and 0.1 were considered to be evolving under relaxed purifying selection [55]. *dN* and *dS* values were also obtained from CODEML. To screen the genes that have undergone positive selection at some sites, two comparisons were performed: M1a vs. M2a and M7 vs. M8, using the constrained (M1a and M7) and unconstrained (M2a and M8) models. Positive selection was further tested using BUSTED.

### Additional analyses of DNA replication and repair genes

33 genes involved in DNA replication and repair were searched against the genome assemblies of seven genera of ancient symbionts (Table S3) downloaded from NCBI, as well as *Walczuchella*. The genome assemblies were annotated using Prokka and BLASTp searches, and their orthologous genes were sorted using OrthoFinder. Further analyses were done for *dnaEQ* and *dnaN* genes of *Walczuchella*, compared to *Karelsulcia, Blattabacterium* and free-living *Flavobacterium*. Gene trees were reconstructed based on the aligned amino acid sequences using IQ-tree with 1,000 bootstrap replicates [56]. Ancestral sequence reconstruction was performed using GRASP [57] to infer insertion and deletion events within the two genes, and the substitution variants were identified through marginal reconstruction in GRASP. Protein structures of *dna* genes were predicted for representative lineages of each symbiont using Phyre2 [58]. Multiple sequence alignments were visualized with the ETE toolkit [59].

### Phylogenetic analyses

The 16S rRNA and 23S rRNA gene sequences of symbionts were extracted using Barrnap v3 (https://github.com/tseemann/barrnap) from metagenome assemblies of giant scales and other bacterial genomes (or individual sequences) available from NCBI. The two genes were aligned individually using MAFFT v7 and concatenated using Geneious Prime. Ambiguous nucleotides were edited using trimAL v.1.4.1 [60] with the *gappyout* option. For the genome-based phylogenetic analysis of *Walczuchella*, amino acid sequences of 134 single-copy orthologous genes were aligned using MAFFT and concatenated into a single matrix with Phyutility v.2.7.1 [61]. The sequence matrix was trimmed using trimAL v1.4.1 with the strict option. For the genome-scale phylogeny of host insects, the near-universal single-copy orthologs (USCOs) were obtained from each metagenome assembly using Patchwork v0.5.1 [62]. The initial set of 2,157 USCOs was recovered from the chromosome-level genome assembly of *Phenacoccus solenopsis* [63] by BUSCO v.5.4.2 [64] utilizing the BUSCO sequences for Hemiptera. Using this initial USCOs set, 1,772–2,097 USCOs were mined from the assemblies of ingroups and outgroups using Patchwork. Protein sequences of the USCOs were aligned with MAFFT and concatenated. The concatenated dataset was trimmed with trimAL v1.4.1 with the *nogaps* option. Maximum likelihood trees for all the above datasets were inferred using IQ-tree under the best-fitting models selected automatically [65]. Branch support was estimated using 1,000 replicates of the ultra-fast bootstrap approximation [66]. The resulting trees were visualized using Figtree v1.4.4 [67].

### Fluorescence *in situ* hybridization

To examine the localization of *Walczuchella* in the host insects, fluorescence *in situ* hybridization was performed on *Crypticerya multicicatrices* and *Icerya purchasi*. The 16S rRNA probe CFB319 [14] that matches *Walczuchella* was used. All insect samples were fixed in 4% paraformaldehyde in phosphate-buffered saline (PBS-1X) and Carnoy’s solution overnight, respectively. The samples were washed with 80% ethanol and bleached with 6% hydrogen peroxide solution [68] for three weeks with several replacements of the solution. They were then washed with absolute ethanol and PBST. Except for the samples to be whole-mounted, the remaining tissues were embedded in paraffin and sectioned to 10 μm with a rotary microtome (HM 340E, Epredia). They were dewaxed with Clear Plus (Falma, Japan) and rehydrated with 100%, 70%, and 50% ethanol series, after drying the sections on slides. The sections were incubated with 300 μl of hybridization buffer containing DAPI (1μg/μl) and probes (100 nM) in a humidity chamber overnight at 45°C. The samples were washed with PBST and mounted with ProLong Diamond Antifade Mountant (Thermo Fisher Scientific), and the tissues were examined under the Nikon Eclipse Ti2-E inverted microscope.

## Results

### Characteristics of *Walczuchella* genomes

All genomes of *Walczuchella* from six host species were closed into circular-mapping chromosomes (Fig. 1A; Table 1). The genome size of *Walczuchella* ranges from 281,644 bp to 309,299 bp, including the largest genome, *Walczuchella* LLAX, previously reported in [27]. The genomes encode 262–320 CDSs, with 12–25 hypothetical proteins. Among the CDSs, 27–86 were predicted to be pseudogenes, mostly due to truncation or fragmentation. All seven *Walczuchella* genomes were completely syntenic (Fig. 1B). The largest proportion of genes in *Walczuchella* genomes was observed in the COG category of information storage and processing (44.5%), followed by metabolism (31.5%) and cellular processes and signaling (12.6%) (Fig. 2B).

**Table 1.** Genome statistics of *Walczuchella* endosymbionts from seven giant scales.

| Symbiont | Genome size (bp) | GC (%) | CDS | Pseudogenized CDS | CDS coding density (%) | rRNA | tRNA |
| --- | --- | --- | --- | --- | --- | --- | --- |
| *Walczuchella* CRMU | 288,630 | 32.4 | 262 | 36 | 79.3 | 3 | 34 |
| *Walczuchella* CRGE | 281,644 | 32.4 | 263 | 27 | 81.5 | 3 | 34 |
| *Walczuchella* DRCO | 295,965 | 32.8 | 313 | 86 | 80.5 | 3 | 34 |
| *Walczuchella* DRPI | 297,114 | 32.8 | 320 | 82 | 80.6 | 3 | 34 |
| *Walczuchella* ICPU | 288,943 | 32.6 | 270 | 32 | 83.4 | 3 | 35 |
| *Walczuchella* ICSE | 288,445 | 32.0 | 282 | 41 | 82.1 | 3 | 34 |
| *Walczuchella* LLAX | 309,299 | 32.7 | 298 | 36 | 86.2 | 3 | 34 |

**Fig. 1.**
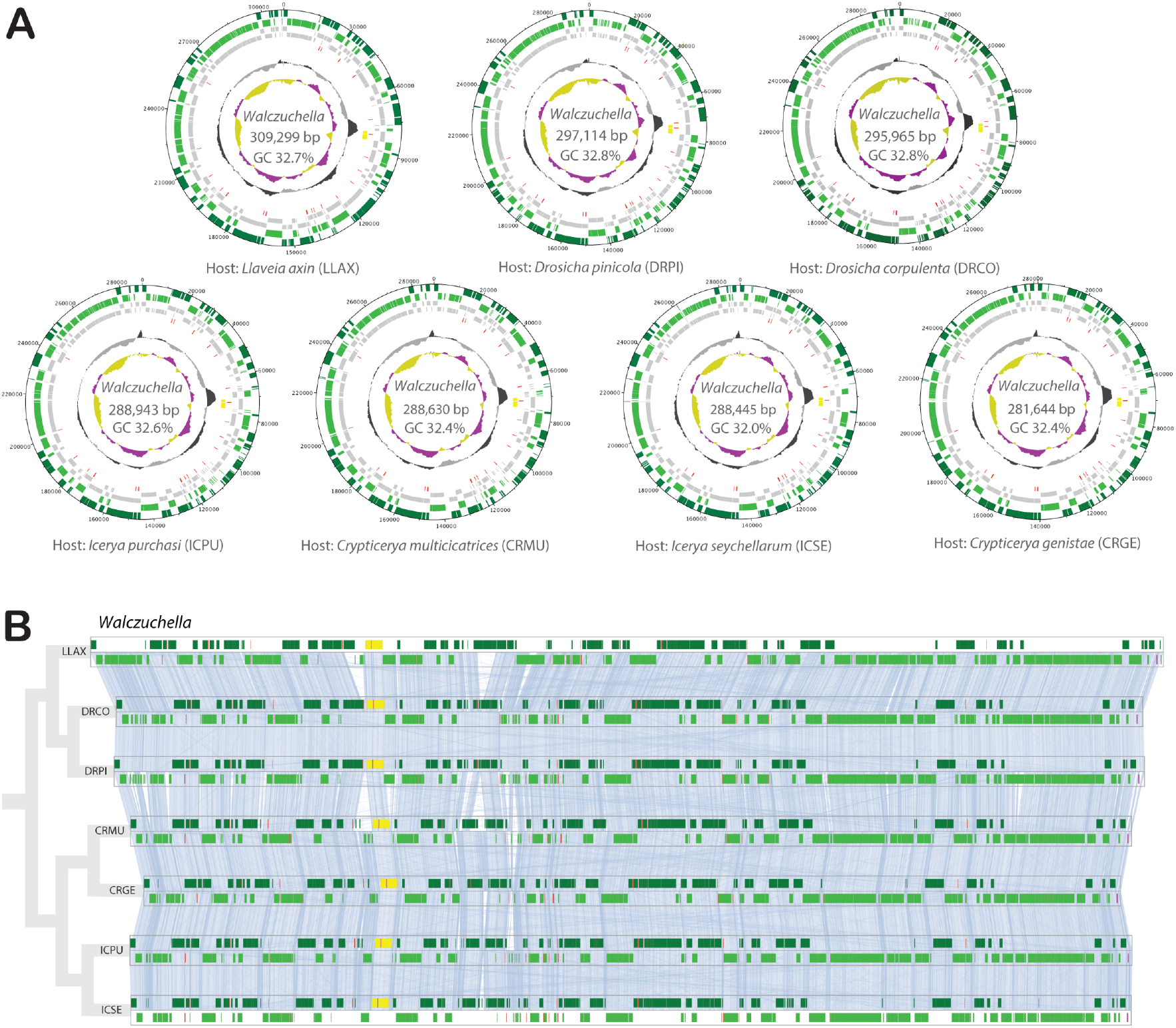
Genome assemblies of *Walczuchella*. **A** Circular genome maps of *Walczuchella*. The lines from outside to inside represent: (i) CDSs on the forward strand (dark green), (ii) CDSs on the reverse strand (light green), (iii) genes on the forward strand (gray), (iv) genes on the reverse strand (gray), (v) tRNA genes (red), (vi) rRNA genes (yellow), (vii) GC plot (above average with black and below average with gray) and (viii) GC skew (above average with purple and below average with yellow). Genome size and GC% are provided in the center of circles. **B** Linear genome alignments of *Walczuchella* with linked lines between colinear genes. *Walczuchella* genomes are collinear without distinct gene rearrangement. Alignments are ordered by the phylogenetic relationship of *Walczuchella* (Fig. S6). The genes are color coded as the genome map above in addition to miscRNA with magenta.

**Fig. 2.**
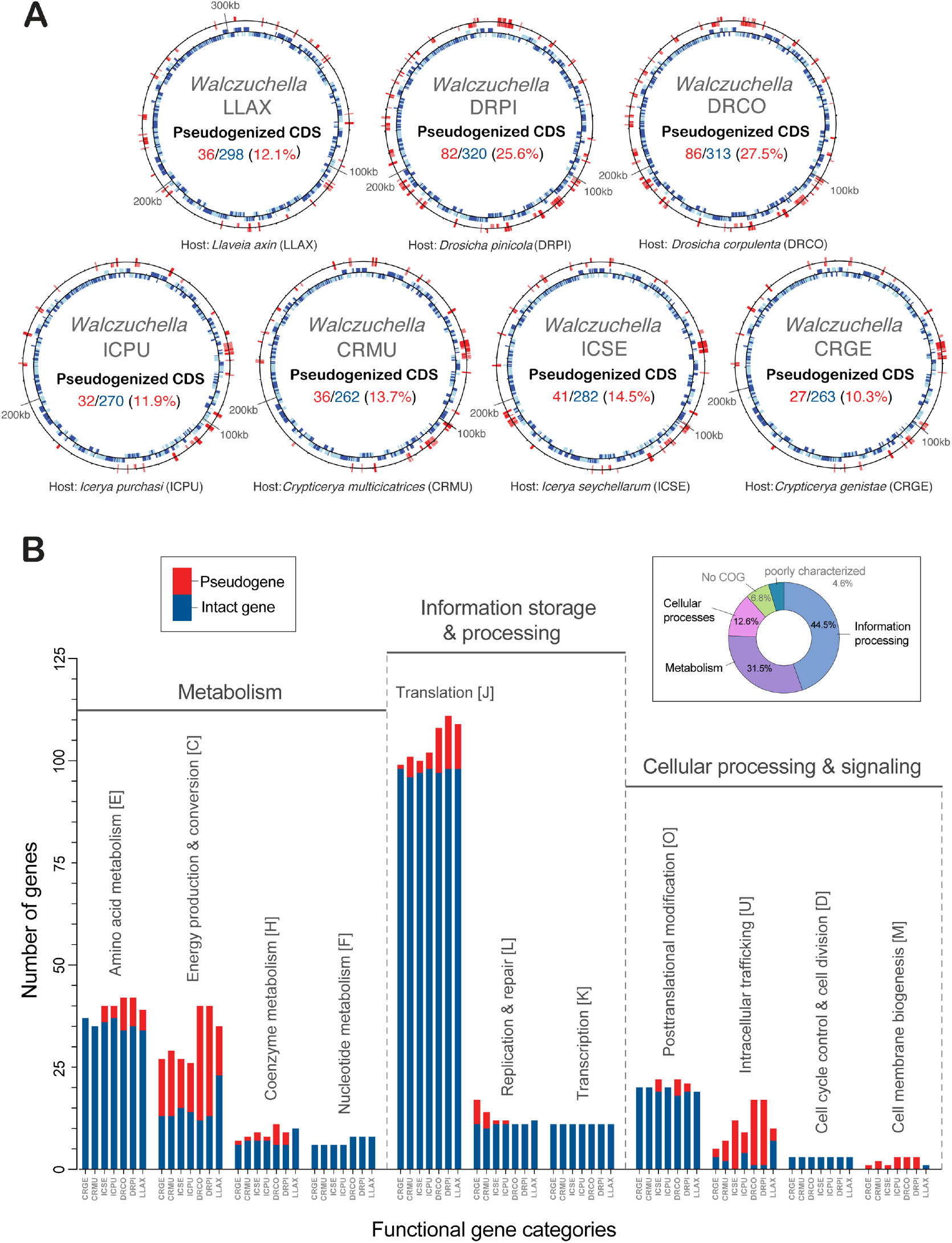
Pseudogenization of *Walczuchella* genomes. **A** Genome maps of *Walczuchella* genomes with pseudogenes (red) and intact genes (blue) on the outer and the inner circles, respectively. **B** Protein-coding genes of *Walczuchella* genomes classified into the clusters of orthologous groups (COG) categories. Intact genes (blue) and pseudogenes (red) are depicted in the bars plot. Average COG proportion of *Walczuchella* genomes is additionally presented right-sided above.

### Gene loss and pseudogenization of *Walczuchella*

The *Walczuchella* genomes showed a high frequency of gene loss and pseudogenization in the following functional COG categories: the biosynthesis of the cellular membrane (M), intracellular trafficking, secretion and vesicular transport (U), and energy production and conversion (C) (Fig. 2; Table S2). In the category (M), the genes *lgt* and *rseP* were only found, but they are pseudogenized or missing in most *Walczuchella* genomes. For the category (U), the majority of 11 genes detected are either lost (in *Walczuchella* DRCO, DRPI, and ICSE) or pseudogenized in the remaining species. In the category (C), out of 38 genes detected in *Walczuchella* genomes, most showed a high frequency of gene loss and pseudogenization, with only a few genes remaining intact. The other COGs showed lower degradation, but we note stochastic losses and pseudogenization of essential genes involved in central informational processes, such as DNA replication and repair, transcription, and translation. Degradation of genes encoding the DNA mismatch repair protein MutS, DNA topoisomerase I, DNA helicase, and endodeoxyribonuclease was observed in the DNA replication and repair category. Missing transcription regulator-encoding genes (*norR* and *glnG*) were also noted in the transcription category. Interestingly, some genes encoding ribosomal proteins of the large subunit were lost or pseudogenized, while the genes encoding the small subunit ribosomal proteins remained intact. Some losses of aminoacyl tRNA synthases were also observed. In the translation category, more than one genome of *Walczuchella* showed gene loss or pseudogenization in genes related to tRNA biosynthesis, ribonuclease, elongation factor 4, peptide chain release factor 1, and translation initiation factors IF-1 and IF-2. Some losses/pseudogenes were also observed in genes responsible for amino acid transport and metabolism.

### Substitution rates and selection pressure on protein-coding genes of *Walczuchella*

The substitution rates of 181 single-copy orthologous genes were estimated to average *dN* = 0.0429 and *dS* = 0.3477 (Figs S1; Table S2). The energy production-related gene *pfkA* had the highest *dN* (0.225) and the second highest *dS* (1.0945), while the 30S ribosomal protein gene *rpsU* had the highest *dS* (1.2213). The results from BUSTED and CODEML showed different averages and ranges for the ω values (Fig. S2; Table S2). The average ω values from BUSTED (0.1670 and 0.1521) were higher than the average from CODEML (0.1340). However, they were congruent in that about 70% of genes (n = 121) showed ω > 0.1, indicating that they were under relaxed purifying selection. The information storage and processing category had the highest number of genes under relaxed purifying selection with 50 genes, followed by the metabolism category with 34 genes, and the cellular processing and signaling category with 12 genes (Fig. 3A; Table S2). Among translation-related genes, six showed relatively high ω values (0.245–0.396; hereafter ω from CODEML), encoding the 50S and 30S ribosomal proteins, a ribosome-recycling factor, and ribonuclease P protein component. In the DNA replication and repair category, the gene *yqeN* had the highest ω value (0.244). In the metabolism category, the genes encoding histidine biosynthesis protein (*hisI*) and anthranilate phosphoribosyl transferase (*trpD*) showed relatively high ω values (0.504 and 0.288, respectively). In the posttranslational modification, tRNA biosynthesis protein TsaB had the highest ω value (0.363). Of the 181 single-copy orthologous genes, 63 may have experienced diversifying selection (Fig. 3B; Table S2). Both analyses of CODEML supported the positive selection of five genes at a *p*-value of 0.01 and 37 genes at a *p*-value of 0.05. In the results of BUSTED, the evidence for the positive selection was significant for six genes at a *p*-value of 0.01 and 9 genes at a *p*-value of 0.05. In all the analyses using CODEML and BUSTED, *hisI, korB, gltX*, and *rpsU* were found to be strongly supported as unconstrained genes.

**Fig. 3.**
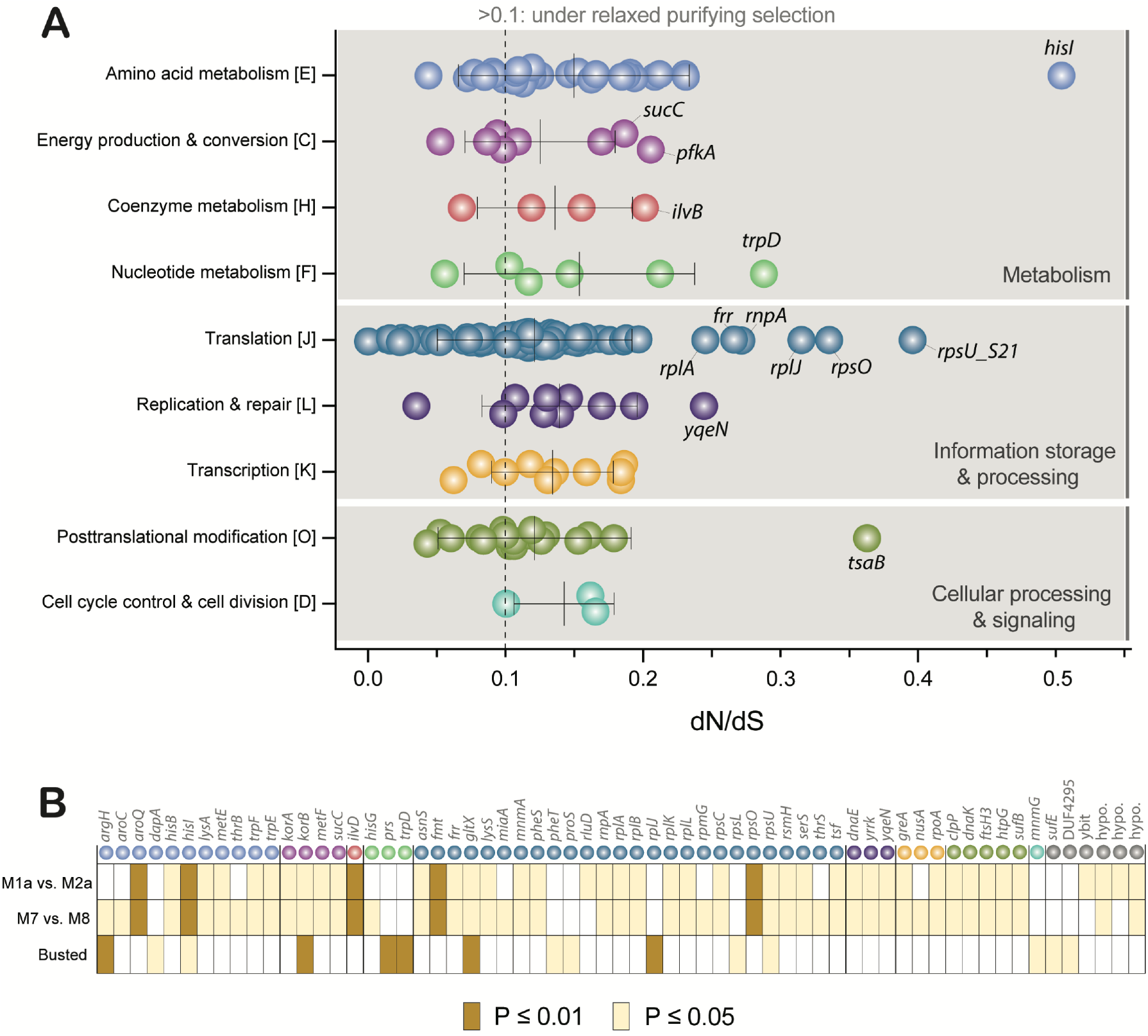
Selective pressures on protein-coding genes of *Walczuchella*. **A** Ratios of non-synonymous to synonymous substitutions (*dN*/*dS*; ω) of single-copy orthologous genes of *Walczuchella*, sorted into the COG categories. Each circle shows ω of the gene estimated using Codeml based on the sequences of seven *Walczuchella*. The names of genes are given around circles showing higher ω in each category. The genes with ω > 0.1 are considered under relaxed purifying selection. **B** 63 genes predicted to have experienced diversifying selection. The first and second rows represent the two results from different approaches (M1a vs. M2a and M7 vs. M8) with Codeml. The third row shows the result of Busted. Each box of genes is filled with different colors following their statistical significance.

### DNA replication and repair genes of *Walczuchella* and other ancient symbionts

The genomes of ancient symbionts were found to contain 33 genes related to DNA replication and repair (Fig. 4; Table S3). DNA polymerase-related genes are relatively conserved across small genomes of all ancient symbionts, in contrast to DNA repair and recombination protein genes *Walczuchella* retains most polymerase genes, but the genes *dnaB* and *dnaG* were absent in all seven genomes. Although *Candidatus* Carsonella and some *Karelsulcia* with smaller genomes lack *dnaB*, this gene is still present in *Candidatus* Nasuia and *Candidatus* Tremblaya, and other ancient symbionts. Among the repair genes, only *mutS* encoding DNA mismatch repair protein is present in three genomes of *Walczuchella*. Genetic properties of *dnaEQ* and *dnaN* genes indicated that mutation accumulation and deletions had occurred in their amino acid sequences, although overall protein structures were found to be similar to those of free-living bacteria (Figs S3, S4).

**Fig. 4.**
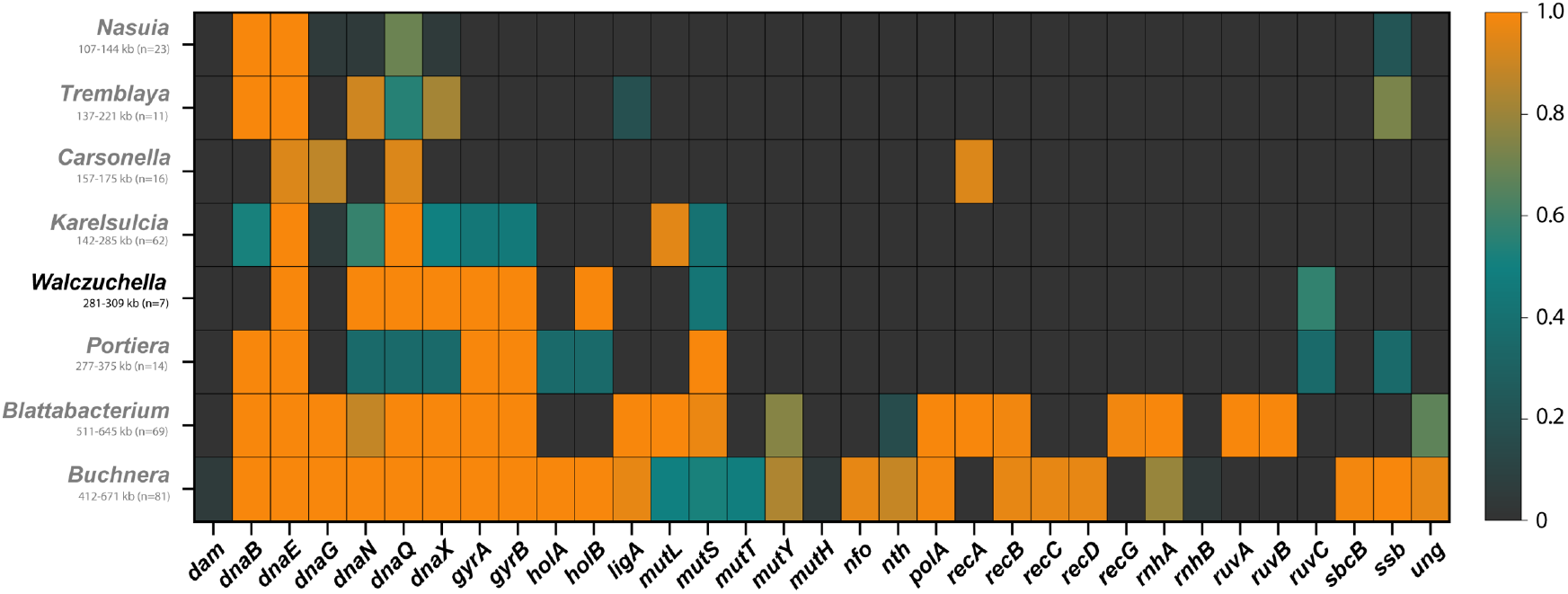
Heatmap showing retention proportions of DNA replication and repair genes in genomes of ancient symbionts. The ranges of genome size and the number of analyzed genomes are noted under the names of symbionts.

### Phylogenetic analyses of *Walczuchella* and their host insects

In the phylogenetic tree based on the 16S and 23S rRNA gene sequences, *Walczuchella* formed a distinct clade comprised of two subclades with *Walczuchella* from the host genera *Drosicha* + *Llaveia* and *Crypticerya* + *Icerya* (Figs. 5A, S5). This topology was confirmed by the phylogenomic analysis based on 134 single-copy genes (Fig. S6A). Additionally, the phylogenenomic tree of host insects, reconstructed using 2,097 USCOs with high bootstrap values, was found to be congruent with the phylogenomic tree of *Walczuchella* (Fig. 5B). The clade of *Walczuchella* was found to be sister to *Candidatus* Hoataupuhia, an obligate symbiont of the Coelostomidiidae (Figs S5). Flavobacterial symbionts of *Cryptococcus* scale insects and *Geopemphigus* aphids were sister to the clade of *Hoataupuhia + Walczuchella*.

**Fig. 5.**
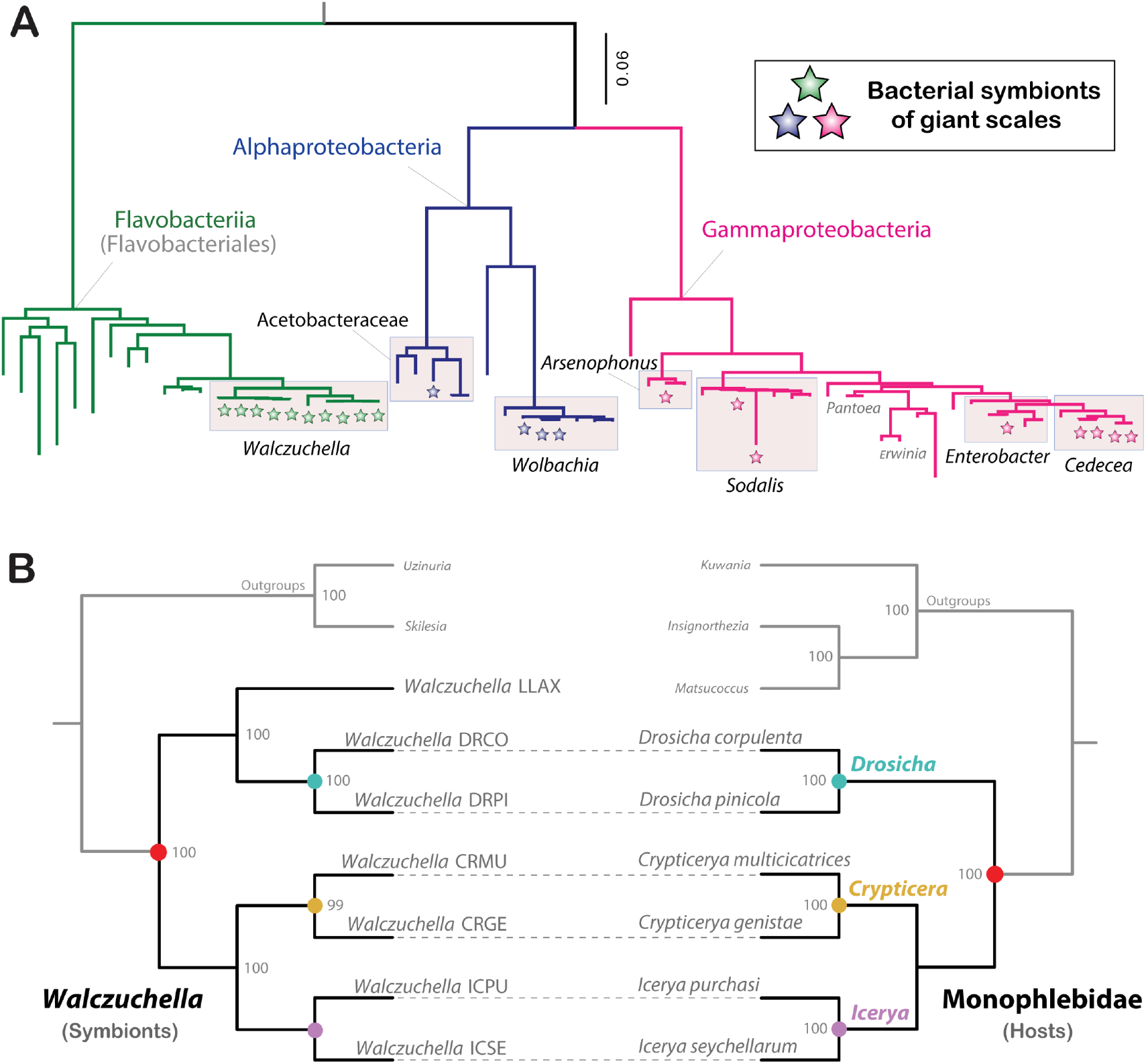
Maximum likelihood analyses of giant scales and their symbionts. **A** Phylogenetic tree of symbionts from giant scales based on 16S-23S rRNA sequences. Symbionts of giant scales are indicated with stars. **B** Co-phylogenetic analysis of *Walczuchella* and their hosts. *Walczuchella* phylogeny (left) is obtained based on 134 single-copy orthologous genes. Host phylogeny (right) is reconstructed based on 2097 USCOs. The numbers at nodes show bootstrap values over 95%. Symbionts are linked with dash lines to their respective host insects. *Walczuchella* LLAX is not linked because of the lack of sequence data of its host. The outgroup taxa are toned down with gray color. All the analyses are inferred with IQ-tree.

### Phylogenetic analysis of co-symbionts

The metagenomes of giant scales revealed the presence of co-symbionts from multiple bacterial lineages, including Alphaproteobacteria and Gammaproteobac-teria (Figs 5A, S5). An Enterobacter-related symbiont was identified in *Icerya purchasi*. Several species of Coelostomidiidae showed *Erwinia*-like symbionts, which cluster close to other Enterobacterales symbionts in giant scales such *Cedecea*- and *Enterobacter*-related lineages. A long-branched *Sodalis*-like symbiont was detected in *Icerya seychellarum. Arsenophonus* was found in *Crypticerya multicicatrices*, and *Wolbachia* was detected in *Drosicha pinicola* and *Icerya purchasi*. Acetobacteraceae-related symbionts were recognized in *Icerya seychellarum* and were phylogenetically close to *Bombella*. No co-symbionts were found in *C. genistae*, which could be due to the sole presence of *Walczuchella* in giant scales or the lower sequencing coverage for this metagenome.

### Localization of *Walczuchella* in giant scales

*Walczuchella* was confirmed in *Crypticerya multicicatrices* and *Icerya purchasi* using FISH and microscopy (Fig. 6). *Walczuchella* signal was localized in the posterior terminal area of the early stage of eggs (Fig. 6A). Its population was divided into two groups in the more developed egg (Fig. 6B) and located on both sides of the posterior area, eventually shaping bacteriomes during embryogenesis (Fig. 6C). The first instar nymph showed clear localization of Walczuchella in about five paired and multilobed structures in the abdomen (Fig. 6D). The localization of Walczuchella in the sectioned tissues was found to be within the bacteriocytes, which had larger polyploid nuclei compared to the normal host cells (Figs 6E-H).

**Fig. 6.**
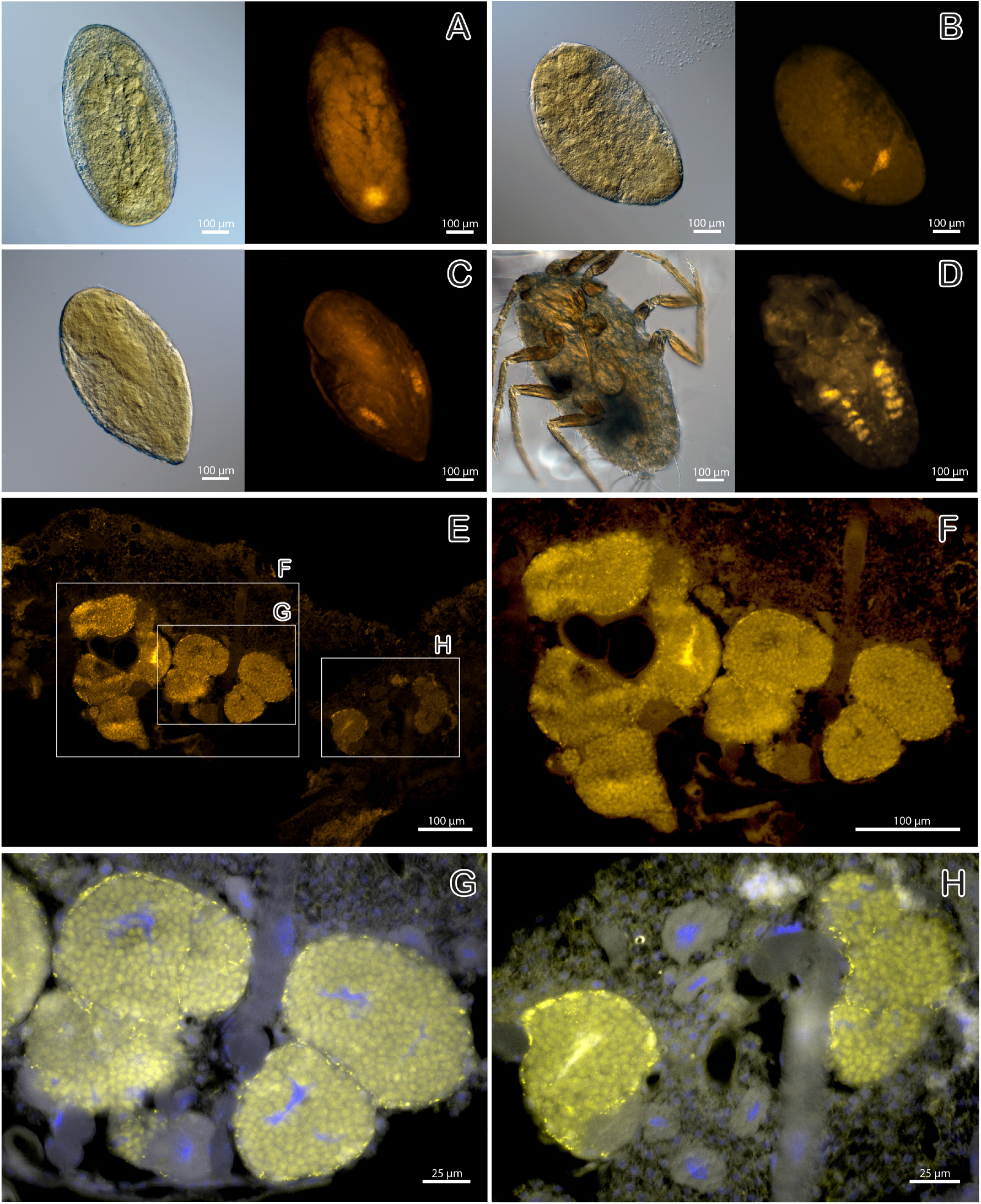
Localization of *Walczuchella* in *Crypticerya multicicatrices* and *Icerya purchasi*. **A** *Walczuchella* is localized in the posterior pore of the early stage of the egg of *I. purchasi*. **B** *Walczuchella* is separated into two groups in the more developing egg of *I. purchasi*. **C** *Walczuchella* is localized on both sides of the posterior part of the egg of *I. purchasi*. **D** Five paired and multilobed bacteriomes of the first instar nymph of *C. multicicatrices* including *Walczuchella*. A-D. Whole mounts. The left photos are light microscopic images, and the right photos are fluorescence in situ hybridization (FISH) images. **E** Entire view of sectioned tissue representing bacteriomes and bacteriocytes of *I. purchasi* in the nymphal stage. **F-H** Higher magnification images of each part in E. E-H. FISH images of body sections. The bacteriomes (roundish structures) contain numerous *Walczuchella* (yellow dots). Nuclei are stained with DAPI (blue) in G and H.

## Discussion

### *Walczuchella* is an ancient symbiont with a large number of pseudogenes

Our results support initial reports from the first sequenced *Walczuchella* genome indicating the ongoing genome erosion in *Walczuchella* [27]. We show that all *Walczuchella* genomes from different host species contain a high proportion of pseudogenes, with up to 27.5% of total CDSs. This is in stark contrast to other ancient symbionts, which usually contain fewer than 5% pseudogenized CDS (Table S4). Furthermore, we show that many intact genes are facing potential deactivation. Approximately 70% of single-copy genes in *Walczuchella* are under relaxed purifying selection (ω > 0.1) (Fig. 3; Table S2). Despite its already highly reduced genome of less than 300 kbp and ∼150 My of coevolution with scale insects, *Walczuchella* seems to be experiencing ongoing and rapid genome erosion. Similar levels of genome erosion are to our knowledge often associated with a rapid shift of the host diet or co-symbiont acquisition. One example of such a change is *Tremblaya princeps* with coding density of 66% in a 138 kbp genome [10]. This genome erosion was likely set in motion by the acquisition of intrabacterial symbionts and subsequent relaxed selection pressure on the genes of *Tremblaya*. However, the reduction of *Walczuchella* is likely more complex as even species with no identified co-symbionts show high numbers of pseudogenes.

### Which functional gene categories are eroding away?

The pseudogenization of *Walczuchella* affects most gene categories, but a few are experiencing extensive erosion (Fig. 2). Significant deterioration was observed in genes associated with the cellular envelope, intracellular trafficking, and energy metabolism (Fig. 2B) which are functions typically lost in tiny endosymbiont genomes due to complementation by the host insect [1]. The genome size of *Walczuchella* between medium-sized and tiny symbionts coincides with losing the capability to synthesize peptidoglycan, phospholipids, and lipopolysaccharides, and produce energy. Transmission electron microscopy of *Icerya* and *Palaeococcus* species showed that *Walczuchella* is surrounded by a host-derived membrane, with no clearly distinguishable bacterial cell wall [69, 70]. A similar cell envelope simplification was, for example, also observed in *Candidatus* Portiera [71, 72]. This phenotype of *Walczuchella*, which is linked to their limited gene set for cell membrane biosynthesis, suggests a large role of the host insect in providing components of the membrane. ATP is provided by the host mitochondria, similarly to other insect symbionts [73] and as suggested by the presence of numerous mitochondria in the bacteriocyte cytoplasm [69, 70] and pseudogenization of *Walczuchella* genes involved in energy production and conversion.

Where *Walczuchella* genomes seem to be more reduced when compared to symbionts with similar genome sizes is DNA replication, transcription, and translation (Fig. 2B). Especially notable are losses of select DNA repair and replication genes (Fig. 4). While other ancient symbionts also experienced substantial losses in DNA repair genes, they usually retain the genes encoding DNA replication proteins. However, *Walczuchella* has lost two key genes, *dnaB* and *dnaG*, which encode the DNA helicase and primase. As with other ancient symbionts, *Walczuchella* has also lost DNA repair genes, although some genomes retain *mutS* encoding the DNA mismatch repair protein. Among translation-related genes, significant decay was found in many aminoacyl-tRNA synthetases and 50S/30S ribosomal proteins. The glutamate-tRNA ligase and 30S ribosomal protein S21 genes were predicted to have undergone positive selection, and several other translation-related genes are under relaxed purifying selection with relatively high ω values (Fig. 3). Other key translation steps such as initiation, elongation, and peptide release (infAB, lepA and prfA) were also affected by the genome erosion. Strikingly, the translational initiation factor IF-1 and some ribosomal proteins genes are missing or present as pseudogenes in some *Walczuchella* genomes even though they are typically retained even in ancient symbionts with tiny genomes [1, 10].

We have confirmed the conservation of genes in the category of amino acid metabolism in the *Walczuchella* genomes (Fig. 2B). This supports its role as a provider of essential amino acids to the insect hosts, as previously reported [27]. However, our findings indicate the loss or pseudogenization of genes encoding enzymes for synthesizing certain amino acids, such as arginine, lysine, methionine, and tryptophan. In addition, the histidine biosynthesis gene (*hisI*) was strongly implicated as having undergone positive selection (Fig. 3). Our analysis suggests that this gene may currently be under relaxed purifying selection, as it showed the highest ω among the genes tested in this study. Rosas-Pérez et al. [27] noted that some substrates for the biosynthesis of several amino acids need to be supplied to complete the pathways due to the loss of amino acid biosynthetic genes (e.g. *aroE, aspC, dapEF, hisCD*, and *ilvE*) in the genome of *Walczuchella*. Our six new *Walczuchella* genomes also confirmed the absence of these genes, while their respective co-symbionts have retained some of them (Table S5). Additionally, horizontally transferred genes of bacterial origin, such as *dapEF*, were found in the host genomes of giant scales (Table S5). These results suggest a complex metabolic patchwork among *Walczuchella*, its co-symbionts, and host genes of both native and bacterial origin similar to other scale insects such as mealybugs [27]. Additionally, we confirm that all *Walczuchella* genomes retain *rpoN*, a regulator of genes involved in nitrogen metabolism and assimilation [15, 27].

### Main drivers of genome erosion in *Walczuchella*

#### Long-term intracellular lifestyle and vertical transmission

*Walczuchella* is a vertically transmitted endosymbiotic bacterium. We confirmed the presence of *Walczuchella* in the eggs of *I. purchasi* and its localization during embryogenesis (Fig. 6A-C). The migration of *Walczuchella* to oocytes was previously observed in the ovarioles of the giant scales *Icerya purchasi* and *Palaeococcus fuscipennis* [69, 70]. In the first instar nymph of *C. multicicatrices, Walczuchella* is present in the paired multilobed symbiotic organ (Fig. 6D), which is similar to the symbiotic organs housing the flavobacterial symbionts of Coelostomidiidae [21, 74]. Phylogenetic analyses reveal that Coelostomidiidae and Monophlebidae, including their symbiotic bacteria, are sister clades (Figs S5) [24, 27, 74]. Furthermore, the symbiotic bacteria and their hosts in Coelostomidiidae and Monophlebidae show co-phylogenetic patterns (Fig. 5B) [21]. These results suggest that they share the most recent common ancestor that originated approximately 125–150 million years ago [24] and their symbionts have coevolved with these two host families since then (Fig. 7B). Similar to many other endosymbionts, the strict intracellular lifestyle and limited population size of *Walczuchella* result in stronger effects of random genetic drift and less effective purifying selection over time [7]. We did not identify any clear differences in these processes between *Walczuchella* and other insect endosymbionts, but we cannot rule out more profound bottlenecking for example due to effective population sizes of giant scales or the number of symbiont cells transmitted from the mother to offspring which are currently unknown.

**Fig. 7.**
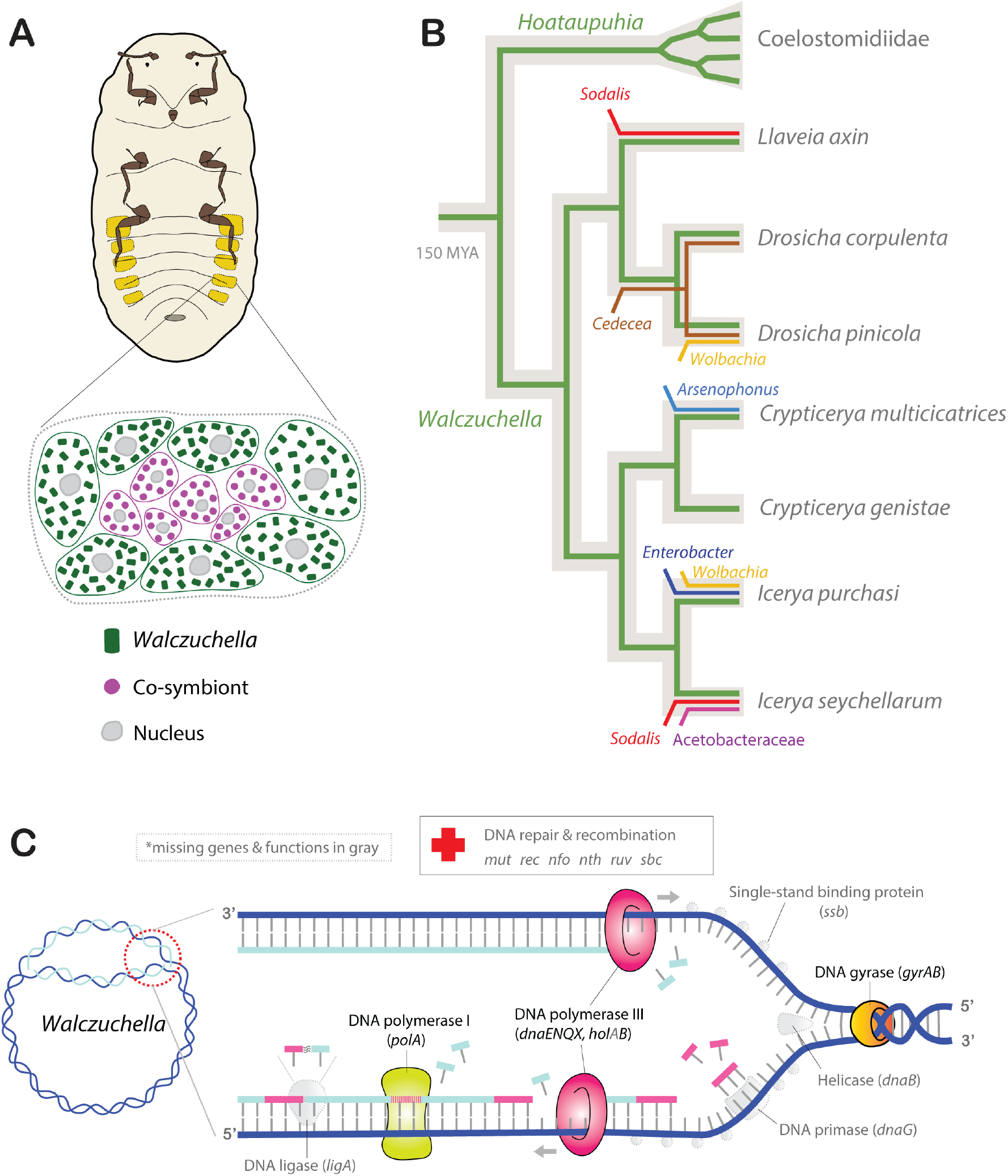
Potential microbial symbiotic features of giant scales leading to the degradation of *Walczuchella* genomes. **A** Intracellular life of *Walczuchella* in bacteriocytes of giant scales with co-symbionts. **B** Long-term evolutionary relationship of *Walczuchella* with hosts and additional introductions of functional symbionts. Colored lines with labels indicate symbionts on host phylogeny (gray background). **C** Incomplete DNA replication and repairing system of *Walczuchella*.

#### Establishment of obligate co-symbionts

The establishment of co-obligate endosymbionts can be an important factor contributing to the ongoing genome reduction of *Walczuchella* in giant scales, except for *Crypticerya genistae* that does not seem to harbor any co-symbionts. We show that various bacterial lineages of Alphaproteobacteria and Gammaproteobacteria are putative obligate co-symbionts of giant scales (Figs 5A, 7B). The composition of these co-symbionts varies depending on the species of giant scales, although *Walczuchella* is uniformly present in all samples. This finding suggests that giant scales have independently acquired diverse co-symbionts during their diversification. *Arsenophonus* and *Sodalis*-allied bacteria are known to complement the missing nutritional genes of ancient symbiotic partners in aphids, mealybugs, and planthoppers [9, 10, 13, 75]. The genome of *Sodalis* TME1 in *Llaveia axin* contains genes involved in all pathways for biosynthesis of essential amino acids, and this *Sodalis* is thus potentially supplying nutrients to the host in cooperation with *Walczuchella* [32]. The acquisition of additional symbionts can be detrimental to *Walczuchella* as more functional ‘junior’ symbionts with a larger gene pool may lead to further relaxation of purifying selection on redundant genes in *Walczuchella*. Cosymbiont acquisition is thus emerging as a major factor driving significant changes in the co-occurring genomes of ancient symbionts that are otherwise relatively stable.

#### Loss of DNA replication and repair genes

The breakdown of the DNA replication and repair system in *Walczuchella* could be a significant factor contributing to their excessive pseudogenization. Our analyses have revealed a limited set of DNA replication and repair genes (Figs 4, 7C). Remarkably, *Walczuchella* has lost two essential genes, *dnaB* and *dnaG*, that are important for the initiation of replication [76]. In comparison, other ancient symbionts, including *Nasuia* and *Tremblaya* with extremely reduced genomes, tend to retain at least *dnaB*. Although there are some cases of *dnaB* loss among *Carsonella* and *Karelsulcia*, their genome size is 157–245 kbp, which is much smaller than that of *Walczuchella* (281–309 kbp) (Table S3). *Sucia* and *Portiera*, which have similar genome sizes to *Walczuchella*, retain *dnaB*. Mutations of *dnaB* were shown to increase genome instability and result in higher mutational frequency in *E. coli* [77].We hypothesize that this early loss of *dnaB* in *Walczuchella* genomes was the tipping point of genome degradation. The remaining DNA polymerase genes have accumulated many mutations likely partly affecting their function (Figs S3, S4). Furthermore, *Walczuchella* does not contain DNA recombination and repair proteins found in symbionts with similarly reduced genomes. The incomplete system of DNA replication and repair leads to an increased rate of mutations that can easily accumulate over time. The enhanced mutation rate is strongly correlated with the evolution of genome size in prokaryotes [4]. The loss of key genes for DNA replication and repair introduces accidental early stop codons, consequently resulting in the more rapid degradation of *Walczuchella* genomes with increased pseudogenization.

### Continuous genome instability of ancient symbionts

Microbial symbionts are subject to environmental conditions that inevitably result in genome reduction, integration with the host, or extinction [78]. The complete genomes of ancient symbionts reveal a wide range of genome sizes, indicating ongoing genome reduction (Fig. S7). Among ancient symbionts of insects, *Buchnera* is a striking example of this phenomenon, with genome sizes ranging from 412 to 671 kbp. *Walczuchella* can be considered an ancient symbiont based on its long-term symbiotic relationship with Monophlebidae and its genome size similar to *Karelsulcia* which originated at least 260 million years ago [14]. Despite its status as an ancient symbiont, *Walczuchella* continues to show unusually high levels of genome degradation. The acceleration is likely due to the early loss of DNA replication/repair genes and the frequent establishment of functional co-symbionts, although we cannot rule out alternative reasons such as unusually strong bottlenecking during vertical transmission, accelerated cellular integration with the host cell, or differences in the host biology such as androdioecy affecting these population genetic processes (Fig. 7). *Walczuchella* joins a few other endosymbionts such as *Tremblaya princeps* in being the exception to the typical pattern of ancient symbionts retaining few pseudogenes [7]. It is a cautionary tale that even if most symbiont genomes may reach a relatively stable state, some such as *Walczuchella* are currently spiraling down the symbiosis rabbit hole of genome instability and pseudogenization. The different rates of genome degradation among symbionts may be due to various factors, such as the mode of transmission, effective population sizes of both hosts and symbionts, differences in host biology, and the genetic bases of mutualistic features [3]. In particular, the number of symbionts transmitted to offspring can be modulated by very specific circumstances as exemplified by *Hodgkinia* in cicadas [79]. Therefore, further work is needed to determine if *Walczuchella* has a lower rate of transmission to offspring compared to other ancient symbionts, as reducing the number of symbionts can promote the fixation of mutations [3].

## Acknowledgments

JYC was supported by the National Research Foundation of Korea (NRF) grant funded by the Korean government (MSIT; 2021R1A6A3A03038909) and JSPS KAKENHI grant (20939772). FH was supported by the JSPS KAKENHI grant (23K14256) and the HFSP Early Career Grant (RGEC29/2024). We thank Nic Schröder for assistance with a phylogenetic analysis of host insects and Dr Yamagishi Kenzo (Meijo University, Nagoya, Japan) for providing us samples of *Drosicha corpulenta* Kuwana. We also acknowledge the Scientific Computing (SCDA), Imaging (IMG), and Sequencing (SQC) sections of the Okinawa Institute of Science and Technology for their great support.

## Author information

JC led the study, conducted the data analyses, prepared all figures, and drafted the manuscript. FH designed the study, supervised the analyses, and revised the manuscript. PP performed DNA extraction and data analysis. HT, TK and MEG provided samples and metagenome data. All authors edited the manuscript.

## Data availability

The genome assemblies of *Walczuchella* and Illumina raw reads are available under NCBI BioProject PRJNA1129605.

---

**Fig. S1.**
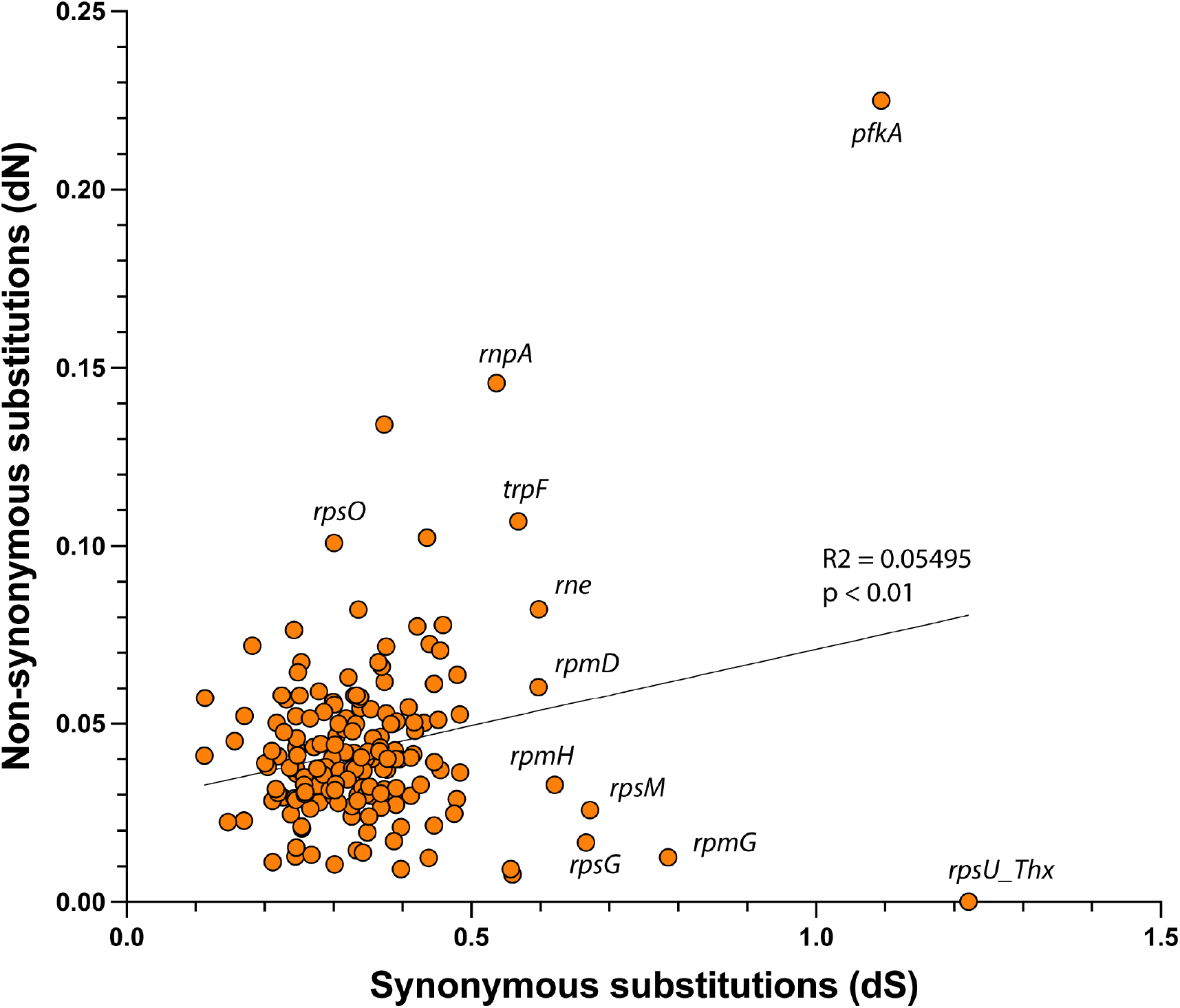
Synonymous (*dS*) and non-synonymous (*dN*) substitution rates of *Walczuchella*. Circles represent the ratio of substitution rates of 181 single-copy orthologous genes of *Walczuchella. dS* and *dN* are estimated using Codeml based on the sequences of seven *Walczuchella*. The gene names are given around circles showing higher substitution rates.

**Fig. S2.**
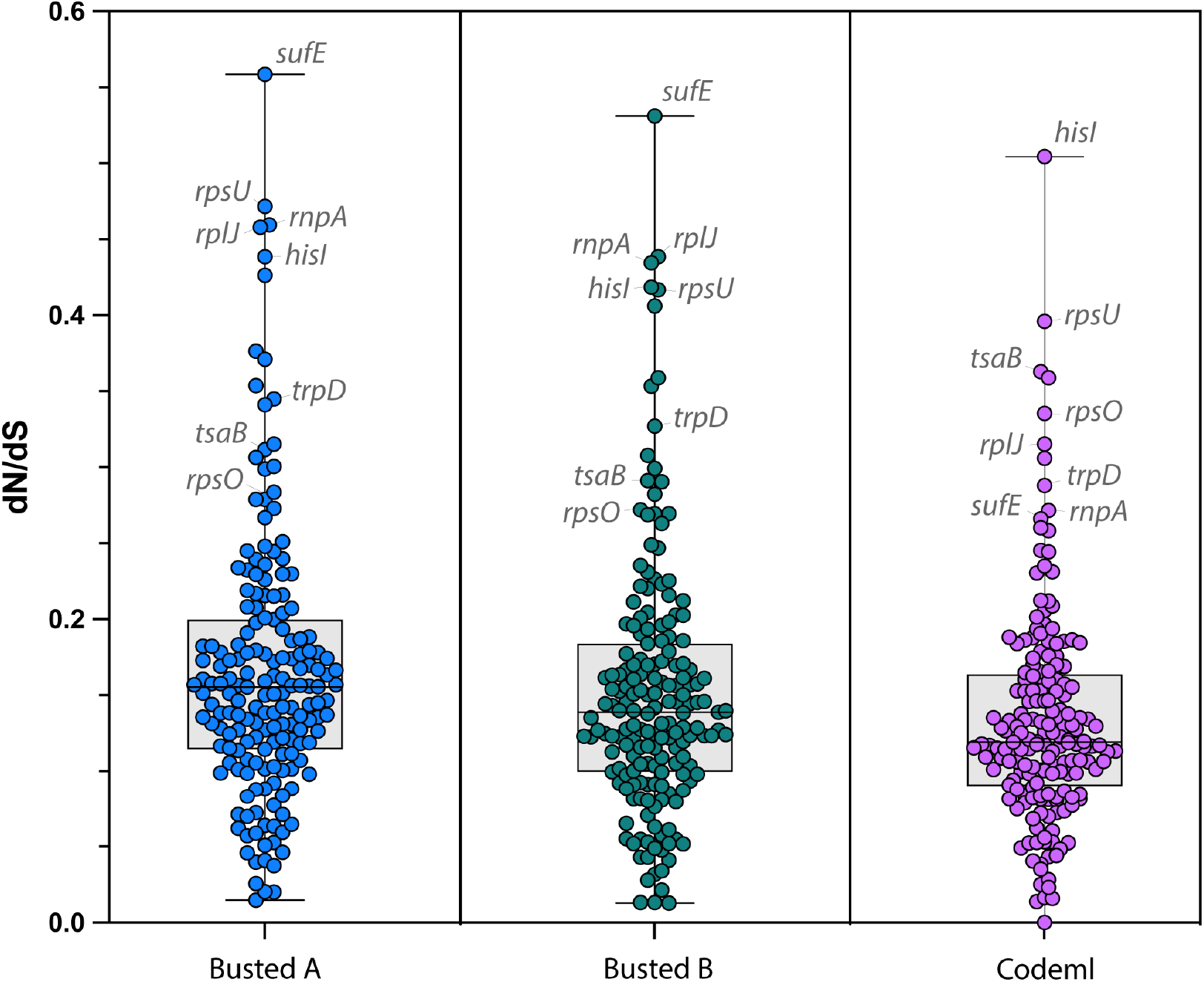
Comparison of ratios of non-synonymous to synonymous substitutions (*dN*/*dS*; ω) estimated using Busted and Codeml. Busted yields two values of ω based on relative GTR branch lengths and nucleotide substitution biases (A), and under a full codon model (B). Each section contains circles based on ω of 181 single-copy orthologous genes of *Walczuchella*. The gene names with higher ω are given around circles for comparison.

**Fig. S3.**
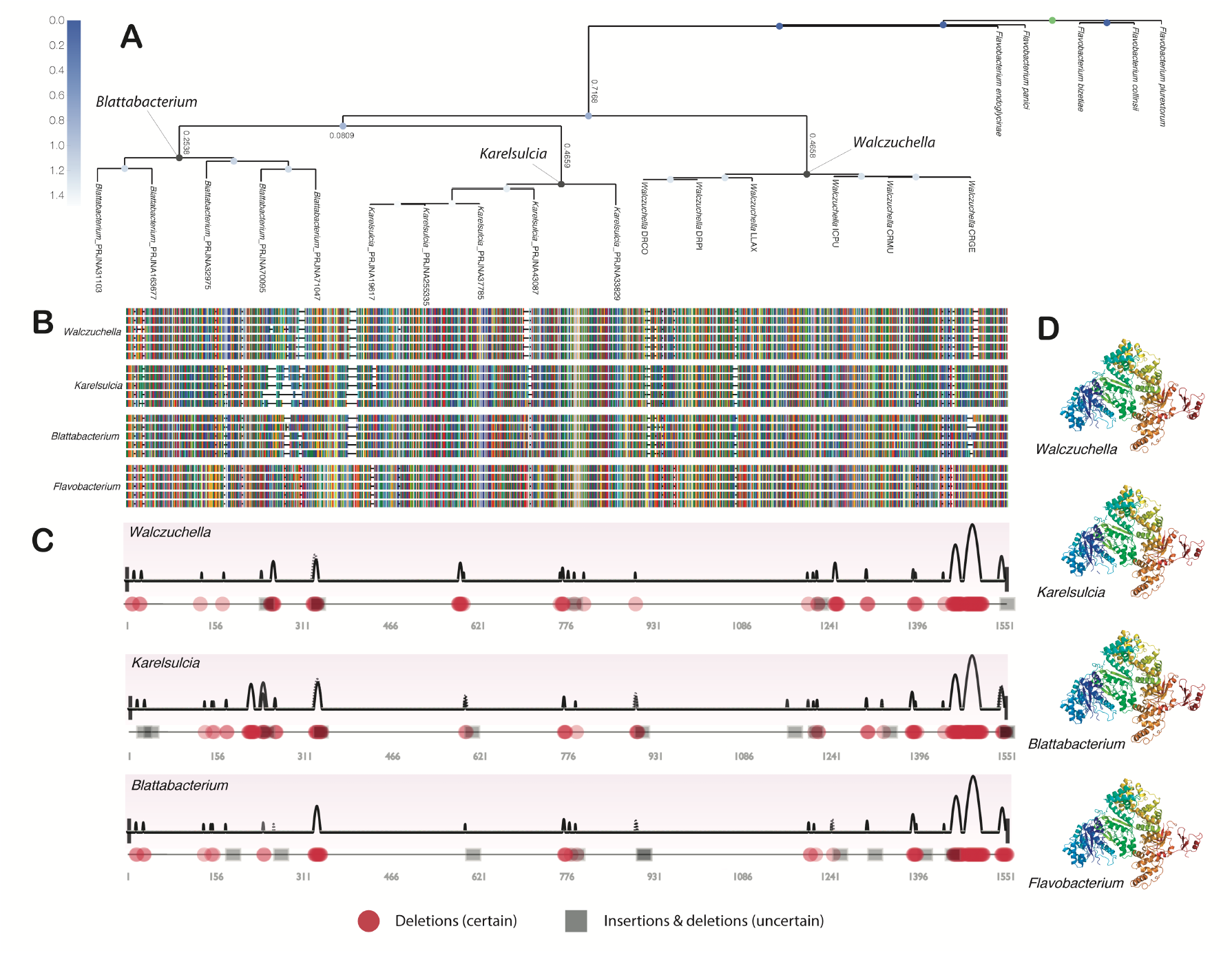
Genetic properties of DNA replication gene (*dnaEQ*) of *Walczuchella*, as well as *Karelsulcia, Blattabacterium* and *Flavobacterium* (free-living). **A** Maximum likelihood phylogeny is inferred with IQ-tree based on 1553 aa sequences of *dnaE*. The branch lengths are present on the main nodes of the tree. **B** Alignment of amino acid sequences of *dnaE*. Each amino acid is shown with different colors and their gaps are filled with dashes. **C** Ancestral sequences of *Walczuchella. Karelsulcia* and *Blattabacterium* are reconstructed with GRASP. The pulse line indicates the inferred mutational regions with strong (solid) and weak (dotted) evidence. Insertions (circle) and deletions (square) are depicted on the respective sites of sequences. **D** Protein structures of *dnaE* are predicted with Phyre2. \***Note**: The phylogenetic analysis of *dnaEQ* (Fig. S3A) revealed that the clade of *Walczuchella* displayed a branch length (0.4658) comparable to that of the *Karelsulcia* clade (0.4659), yet both branch lengths were significantly greater than that of the *Blattabacterium* clade (0.2538). The *dnaEQ* amino acid sequences of *Walczuchella, Karelsulcia* and *Blattabacterium* showed several deletions when aligned with sequences of free-living bacteria (Fig. S3B). The ancestral reconstruction of the *dnaEQ* amino acid sequence showed that *Walczuchella* had experienced 119 deletion events in its ancestral sequence, which was higher than *Blattabacterium* (105) but lower than *Karelsulcia* (137) (Fig. S3C). Despite the accumulated mutations in all three ancient symbionts, their *dnaEQ* protein structures were found to be similar to those of free-living bacteria (Fig. S3D).

**Fig. S4.**
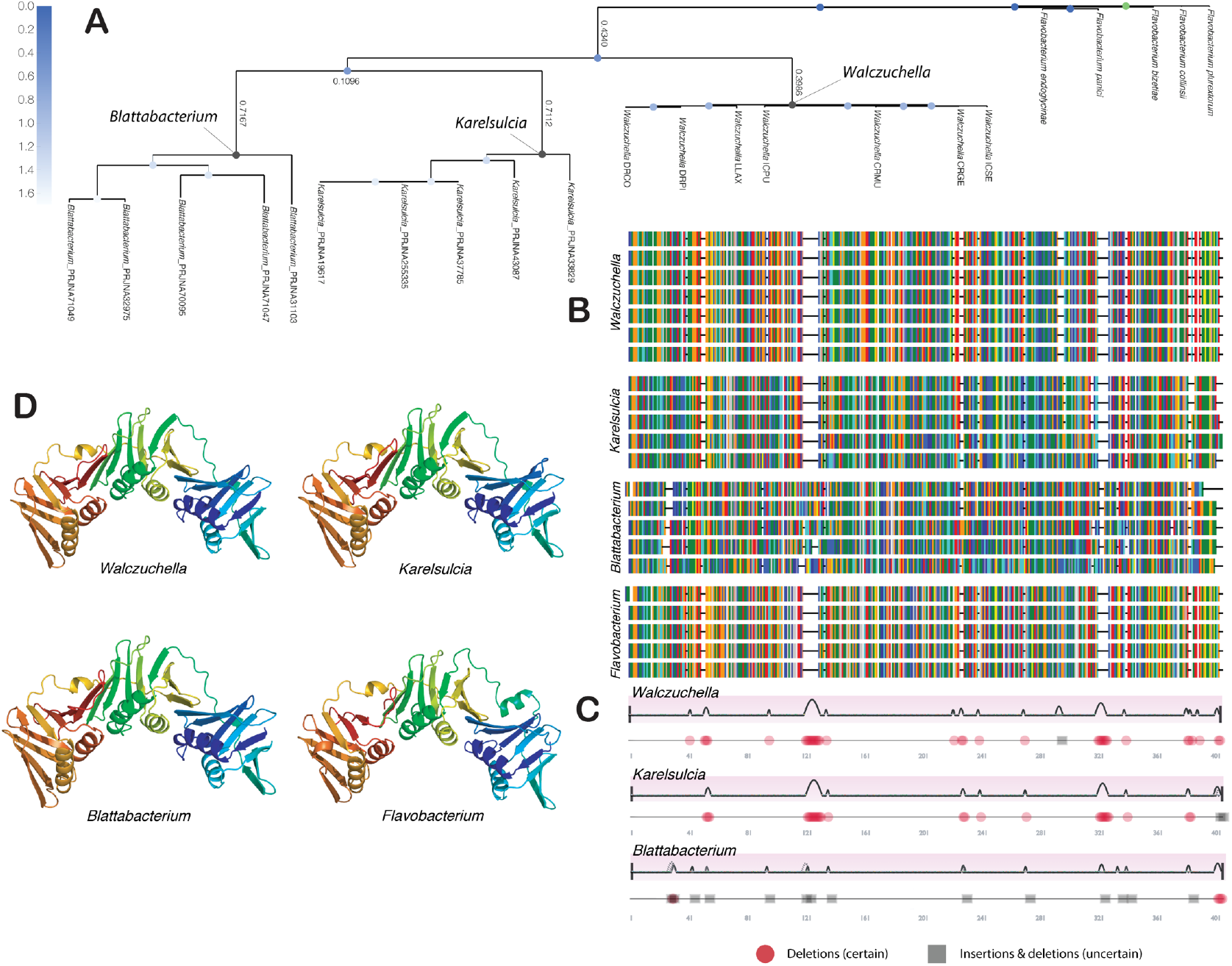
Genetic properties of DNA replication gene (*dnaN*) of *Walczuchella*, as well as *Karelsulcia, Blattabacterium* and *Flavobacterium* (free-living). **A** Maximum likelihood phylogeny is inferred with IQ-tree based on 403 aa sequences of *dnaN*. The branch lengths are present on the main nodes of the tree. **B** Alignment of amino acid sequences of *dnaN*. Each amino acid is shown with different colors and their gaps are filled with dashes. **C** Ancestral sequences of *Walczuchella. Karelsulcia* and *Blattabacterium* are reconstructed with GRASP. The pulse line indicates the inferred mutational regions with strong (solid) and weak (dotted) evidence. Insertions (circle) and deletions (square) are depicted on the respective sites of sequences. **D** Protein structures of *dnaN* are predicted with Phyre2. \***Note**: The phylogenetic analysis of *dnaN* (Fig. S4A) showed that the *Walczuchella* clade had a lower branch length (0.3986) than those of *Karelsulcia* (0.7112) and *Blattabacterium* (0.7167). The *dnaN* amino acid sequence alignments of *Walczuchella* and *Karelsulcia* were similar to free-living bacteria, while *Blattabacterium* had a different alignment pattern resulting from distinct insertions and deletions to the others (Fig. S4B). Ancestral sequence reconstruction of *dnaN* revealed that *Walczuchella* had experienced 39 deletions in its ancestral sequence, which was higher than those of *Karelsulcia* (32) and *Blattabacterium* (18) (Fig. S4C). This indicates that substitutions have likely occurred more often in *dnaN* of *Blattabacterium* and *Karelsulcia* than in *Walczuchella*, but deletions happened more frequently in the gene of *Walczuchella*. The structures of the *dnaN* protein of ancient symbionts and free-living bacteria were found to be similar (Fig. S4D).

**Fig. S5.**
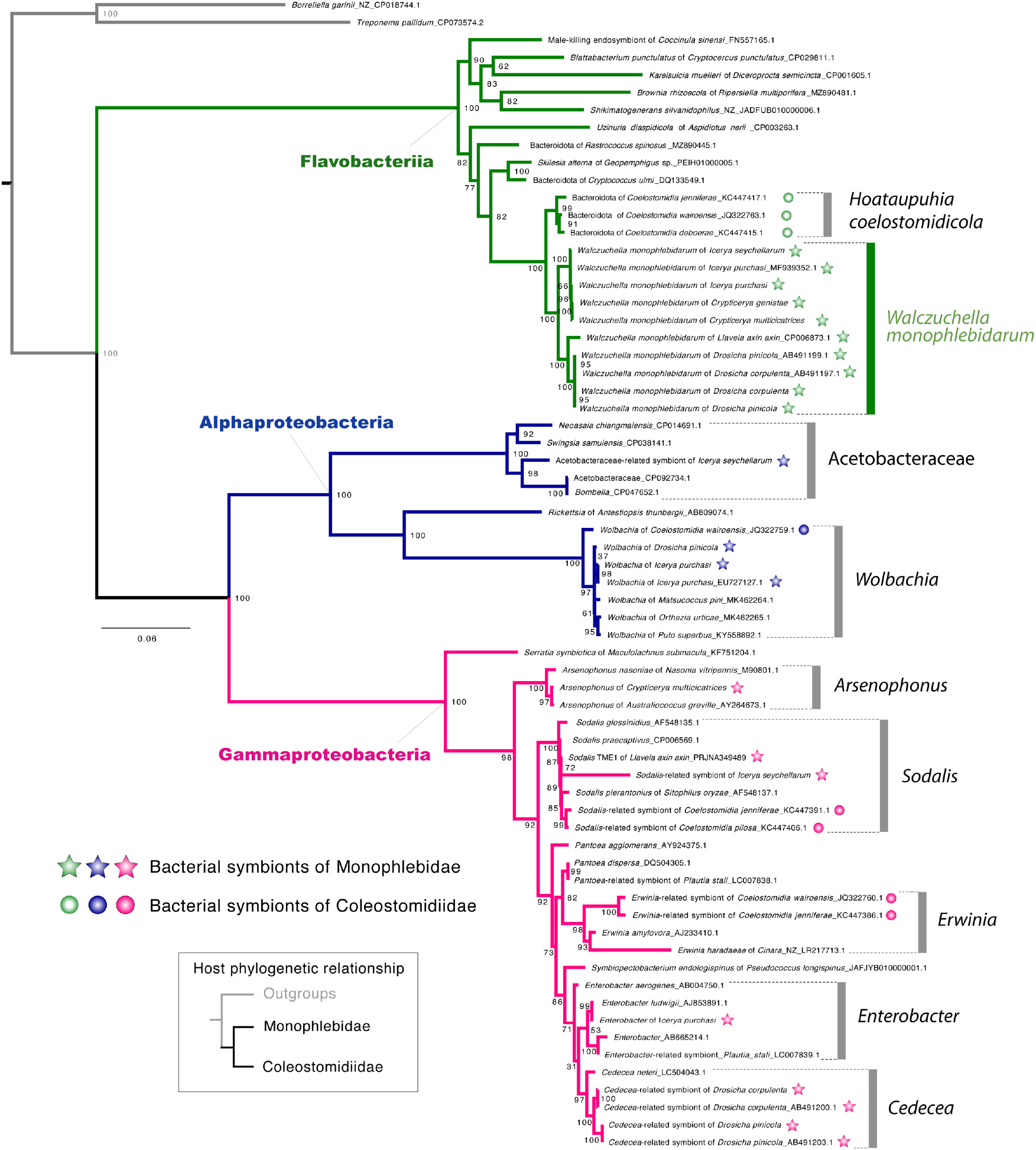
Maximum likelihood analysis of symbionts from giant scales. The phylogenetic tree is inferred with IQ-tree based on 16S-23S rRNA sequences under GTR+F+I+G4 model. Symbionts of Monophlebidae and Coleostomidiidae are indicated with stars and circles, respectively.

**Fig. S6.**
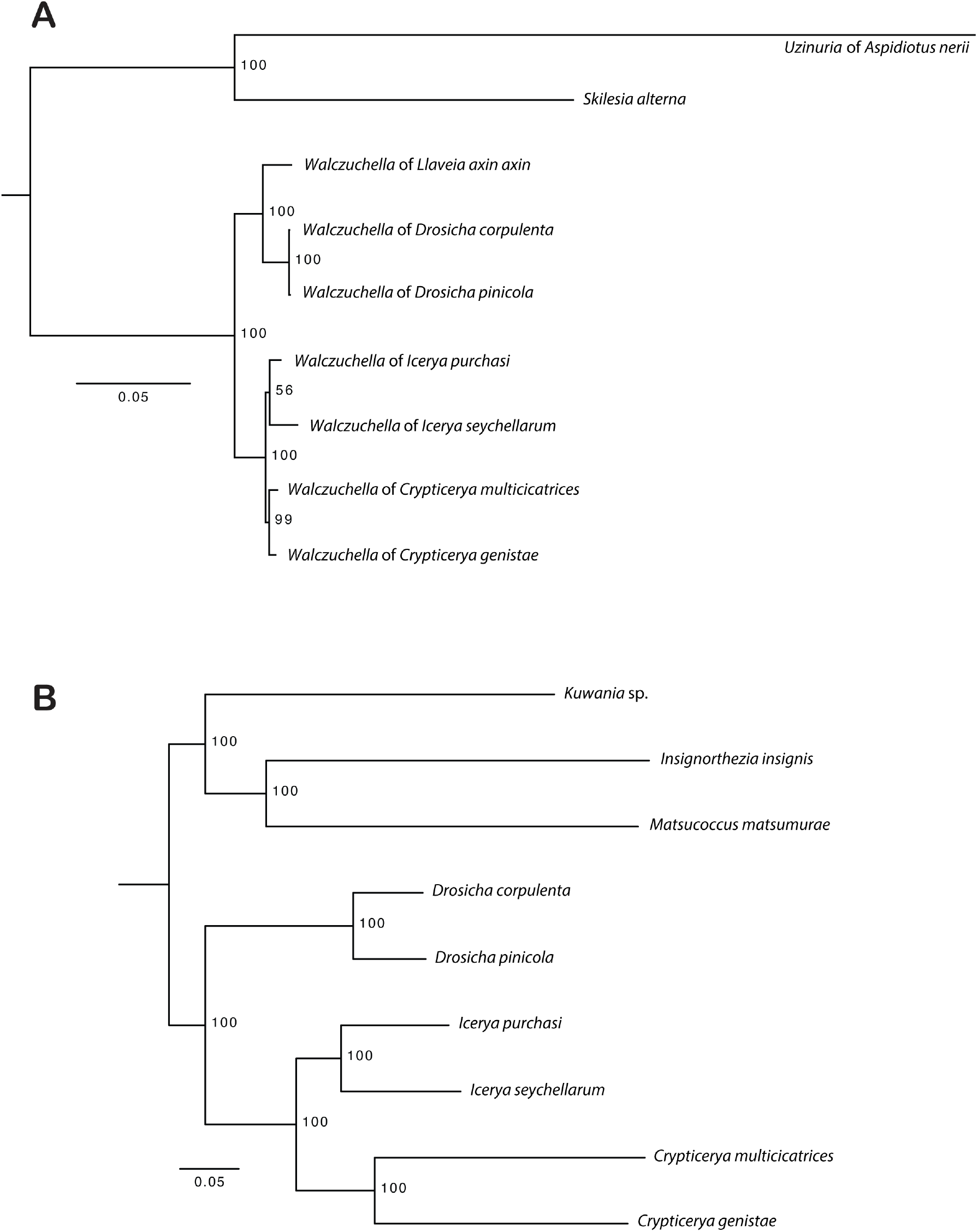
Maximum likelihood analyses of *Walczuchella* and their hosts. **A** *Walczuchella* phylogeny obtained based on 134 single-copy orthologous genes under cpREV+F+G4 model. **B** Host phylogeny reconstructed based on 2097 USCOs under VT+F+I+I+R3 model. All the analyses are inferred with IQ-tree. The numbers at nodes show bootstrap values.

**Fig. S7.**
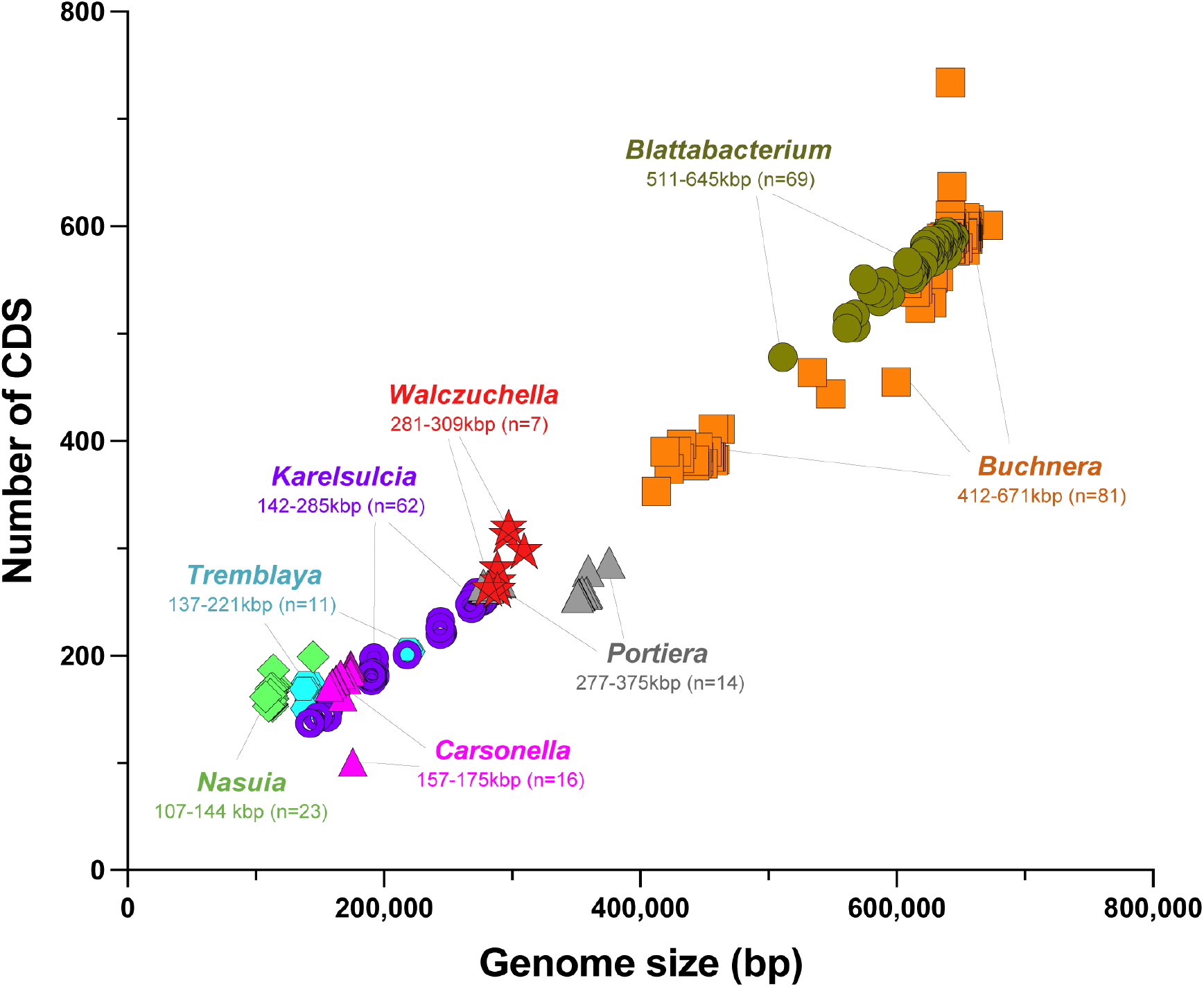
Genome size and the number of CDS of ancient symbionts. The ranges of genome size and the number of genome assemblies used in this plot are noted under the names of ancient symbionts.

